# Direct detection of alternative DNA conformations with long-read sequencing and machine learning approaches

**DOI:** 10.64898/2026.07.17.739277

**Authors:** Jacob P. Sieg, Huqing Zeng, Lauren Heaverly, Kateryna D. Makova

## Abstract

Progress has been made in identifying G-quadruplexes (G4s) and other non-canonical (non-B) DNA structures in live cells. However, these experiments have been limited by methodological constraints, including low resolution and specificity, and GC-sequencing bias inherent to the short-read sequencing technologies. Direct, single-molecule, and long-read technologies have the potential to address these shortcomings. Here, we investigated the use of long-read DNA sequencing with Oxford Nanopore Technologies (ONT) to detect G4 and other non-B DNA structures. We applied ONT sequencing to oligos known to form G4s *in vitro*, as well as to DNA from live cells. Both experiments were probed with potassium permanganate, a chemical that preferentially oxidizes single-stranded DNA (ssDNA). We determined that G4 structures can be resolved by ONT sequencing, and a G4 motif’s sequencing error profile depends on the original structural state of the G4 (as determined *in vitro*). We next applied machine learning algorithms (logistic regression, random forest, XGBoost, and 1D convolutional neural networks) to the raw ONT sequencing current data. Using these approaches, we could determine whether a sequencing read originated from a G4 motif in the G4 vs. B-form conformation with ∼90% accuracy. Further, we demonstrated that it is possible to sequence permanganate-modified DNA directly using ONT, with minimal effects on the read length, yield, and alignment accuracy. We conclude that at high ONT sequencing depth, this approach can identify B-folded vs. single-stranded DNA regions in cells, with ssDNA frequently present across several non-B DNA structures.

## Introduction

Recent experiments have identified G-quadruplexes (G4s) and other non-canonical (non-B) DNA structures in live cells (later called “*in cell*”), revealing functions in gene regulation, genome organization, and chromatin structure (reviewed in ^1,2^). However, current progress in this area is limited in part by constraints imposed by short-read sequencing techniques. Short-read sequencing techniques have low mapping rates in repetitive genomic regions^3^ and substantial bias against GC-rich and non-B-motif-containing sequences^4^. Moreover, current approaches usually require purification of non-B structures, followed by amplification, and cannot report on more than one structure in a single read, thus obscuring the resulting information on underlying populations of structured molecules.

Long-read sequencing techniques have solved some of these problems for other applications. Highly accurate Pacific Biosciences (PacBio) reads, combined with extremely long Oxford Nanopore Technologies (ONT) reads, have resolved complex repetitive regions such as human centromeres and facilitated the assembly of telomere-to-telomere genomes (reviewed in ^5^). Of particular note, GC-bias-tolerant ONT reads enabled assembly of G4-rich dot chromosomes in songbirds for the first time^6,7^. Additionally, the long-read techniques Fiber-seq and DiMeLo-seq provide information on chromatin accessibility and protein-DNA interactions, respectively, at single-haplotype and single-molecule resolution^8,9^.

A key driver of these advances is PacBio and ONT direct DNA sequencing, which enables identification of DNA base modifications in the raw sequencing signal ^10^. This signal consists of the interpulse duration time for PacBio and electrical current features for ONT. For DNA, both technologies currently support calling 5-methyl-deoxycytosine, 5-hydroxymethyl-deoxycytosine, and N6-methyl-deoxyadenosine modifications^10^, which can occur naturally in a sample or be introduced by experimental manipulations. For example, Fiber-seq relies on the simultaneous detection of natural 5-methyl-deoxycytosine and artificial N6-methyl-deoxyadenosine introduced by a DNA methyltransferase to generate a profile of a genome epigenetic landscape^8^.

Similar approaches have been employed to detect DNA structure. Non-B motifs have been shown to slow down PacBio polymerization rates and affect nanopore translocation times^11,12^. However, the ground truth was uncertain in both studies; therefore, it is unknown whether these non-B DNA sequencing signatures were caused by preexisting non-B structure or were inherent to the motifs.

Here, we investigated two strategies for detecting G4 and other non-B DNA structures with ONT. In the first, we pre-folded G4 motifs into a G4 or a B-DNA structure, then sequenced them to identify what sequencing features originate from the G4 structure or the DNA motif. Second, we used chemical reactivity of potassium permanganate to tag ssDNA, which is then read out from raw pore current signals to detect non-B DNA structure via the ssDNA it induces.

## Results

### Nanopore sequencing errors are sensitive to secondary structure at G4 motifs

We first distinguished sequencing errors arising from G4 secondary structure vs. those associated with a B-form helix using a G4-duplex model system *in vitro* (Figure 1A). This G4-duplex model system was previously used to study G4 structure in a duplex DNA context^13,14^. In this system, a G4 motif was placed between two B-DNA helices. The G4 structure was activated by providing an imperfect, poly-T reverse complement (RC), eliminating base pairs (bp) that compete with the G4 structure. Conversely, the G4 motif was forced into a B-form helix by denaturing and annealing in the presence of a perfect RC. Thus, sequencing errors can be attributed to G4 structure or to the motif in a B-DNA form by sequencing in the presence of a poly-T RC or a perfect RC, respectively.

**Figure 1.**
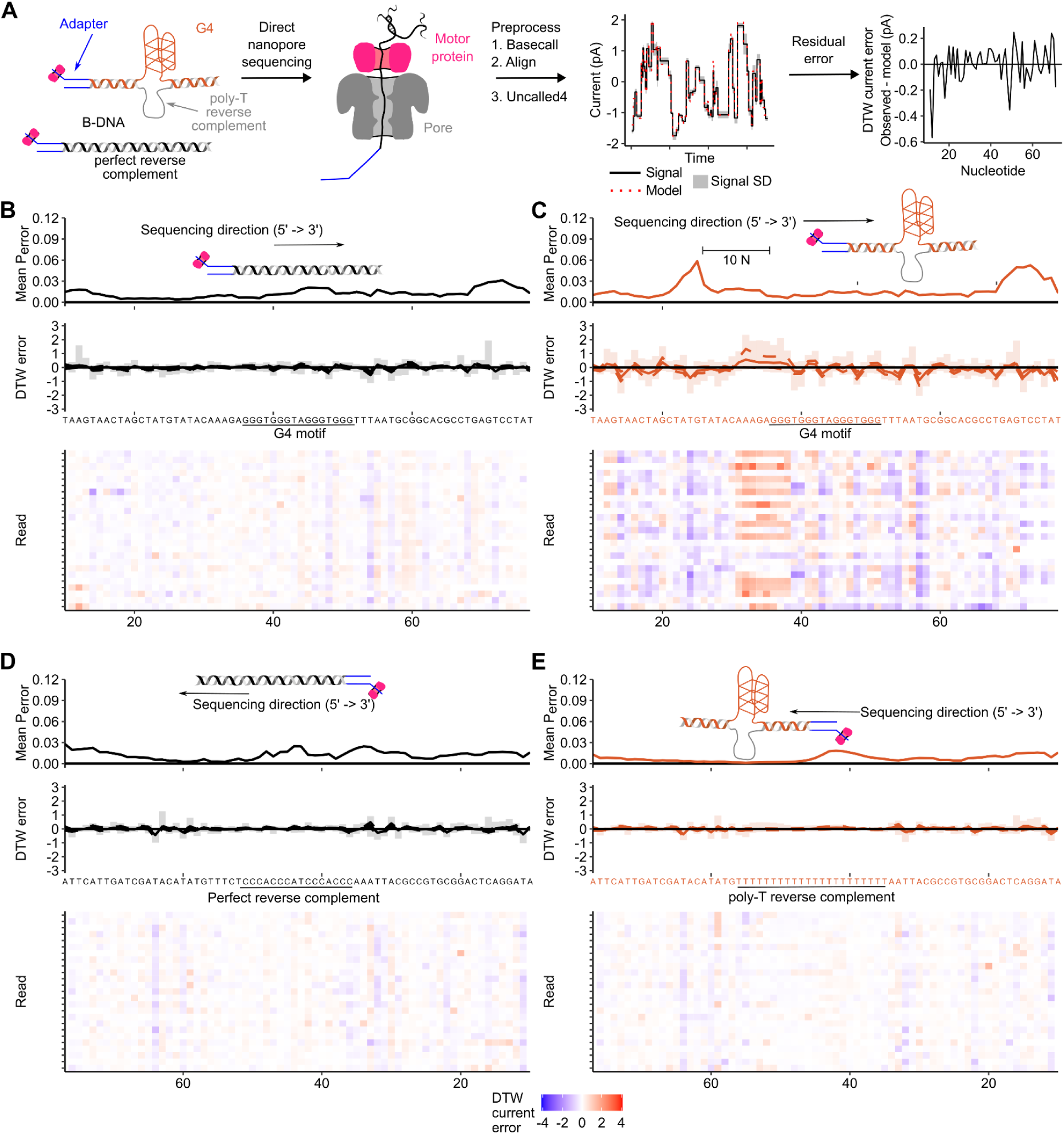
G-quadruplex (G4) structure affects Oxford Nanopore Technologies (ONT) sequencing signal for the *cMYC* promoter G4-duplex model. **(A)** Detection of residual errors in ONT data. Model G4 motifs were folded into a B-form or into a G4 structure and sequenced. Deviations from the pore model were assigned with Uncalled4 software. **(B)** Analysis of error for G4-motif in a B-form helix. X-axis is shared and corresponds to position. Top: Mean probability of a basecall error (P_error_) calculated from Phred scores. Middle: DTW current error in pico amps (pA). Bars, dotted lines, and solid lines represent 95% confidence intervals, 50% confidence intervals, and means, respectively. Bottom: DTW current error heat maps for 25 randomly selected sequencing reads. **(C)** The same plots for the G4 conformation. (**D)** and **(E)** correspond to the same plots for the reverse complement in a B-DNA and G4 conformation, respectively.

We modified three previously characterized G4-duplex systems for ONT sequencing, extending the duplex ends 20 nucleotides (nt) to enable multiplexing and increase the distance between the duplex end and the G4 motif (Table 1). We named these constructs cMYC, VEGF-NMR, and VEGF-WT, based on G4 structures identified previously in the corresponding human genes. The 3D structure of the cMYC G4-duplex model was solved to 7.4 Å, and is derived from the promoter of the human *cMYC* proto-oncogene ^13,15^. The *VEGF* G4-duplex models were used to characterize the effects of DNA damage G4 folding dynamics, chosen because it is a well-characterized site where DNA damage affects the regulation of the human *VEGF* proto-oncogene ^14^. The NMR model was modified relative to the WT, native human G4 sequence, to make the sequence experimentally tractable *in vitro* ^16^. The G4 folding and duplex folding stoichiometry were confirmed using circular dichroism (CD) and native gel electrophoresis, respectively, before ONT sequencing (Figure S1).

**Table 1.**
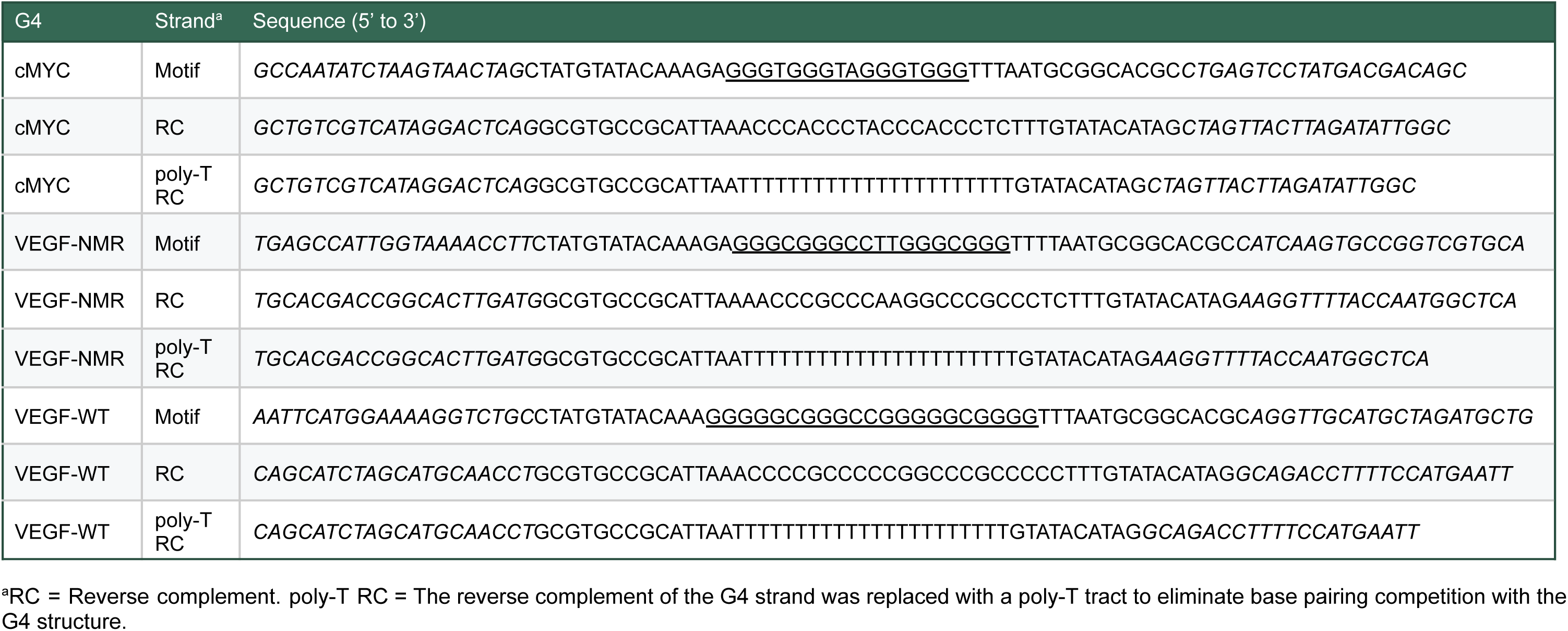
G4 model oligo design. G4 motifs and 20 nt B-DNA extensions are designated by underlined and italic text, respectively.

We next compared sequencing errors and current profiles between G4 structures and B DNA for these three G4-duplex models. Table S1 provides summary statistics for ONT sequencing and alignment of G4 models folded *in vitro*. Data were analyzed with the NanoPrint toolkit, a custom Snakemake pipeline that performs basecalling, alignment, and sequencing error analysis (Figure 1A). The Nanoprint toolkit performs two types of error analysis. In the first, it computes average basecall error probability (P_error_) calculated from Phred quality scores provided by the basecaller (Figure 1B-E, top two panels). The second error analysis compares the pore model to the raw nanopore current signal calculated using the Uncalled4^17^ program (Figure 1B-E, bottom two panels). Uncalled4 uses a basecaller-guided dynamic time warping (DTW) algorithm to accurately align nanopore signals to a reference genome, thus providing DTW current error in pico amps (pA), an information-rich deviation between the pore model and observed nanopore current signal.

Our analysis revealed significantly different error and current profiles for the *cMYC* G4 motif in the B-form helix vs. G4 structure with an effect size of 6.4±0.5 pA^2^ (Figure 1B-C, Table 2, *p* < 0.001 permutation test with subsampling^18^ see Methods for details). For the G4 structure, base call error probability peaked 10 nt to the 5’ end of the G4 motif (Figure 1C top). Likewise, the G4 structure exhibited strong negative DTW current error 10 nt to the 5’ end of the motif, transitioning to strong positive error 5 nt to the 5’ end of the motif (Figure 1C middle). Then the DTW current error was positive in G-run stems and negative in loops (Figure 1C middle). The same analysis was performed for the RCs (Figure 1D and Figure 1E), where there was a significant difference but with a much smaller effect size of 0.20±0.02 pA^2^ (Figure 1D-E, Table 2; *p* = 0.008±0.006). The same features were observed for the VEGF-NMR and VEGF-WT constructs (Figure S2-3, Table 2). In summary, sequencing errors were sensitive to secondary structure on the G4-motif-containing strand but not on the RC strand.

**Table 2.** Results of the permutation test comparing DTW current error profiles between conditions. Full parameters are available in Table S2.

| motif | strand | treatment 1 <sup>a</sup> | treatment 2 <sup>a</sup> | conformation 1 | conformation 2 | median *p*<br>value <sup>b</sup> | *p* value<br>sd <sup>c</sup> | Effect size<br>mean (pA<sup>2</sup>) <sup>d</sup> | Effect size<br>sd (pA<sup>2</sup>) <sup>d</sup> |
| --- | --- | --- | --- | --- | --- | --- | --- | --- | --- |
| cMYC | motif | 0 mM CTRL | 0 mM CTRL | B-DNA | G4 | 0.000 | 0.000 | 6.38 | 0.48 |
| cMYC | RC | 0 mM CTRL | 0 mM CTRL | B-DNA | G4 | 0.008 | 0.007 | 0.20 | 0.02 |
| VEGF-NMR | motif | 0 mM CTRL | 0 mM CTRL | B-DNA | G4 | 0.000 | 0.000 | 6.31 | 0.92 |
| VEGF-NMR | RC | 0 mM CTRL | 0 mM CTRL | B-DNA | G4 | 0.000 | 0.000 | 0.70 | 0.13 |
| VEGF-WT | motif | 0 mM CTRL | 0 mM CTRL | B-DNA | G4 | 0.000 | 0.000 | 3.18 | 0.43 |
| VEGF-WT | RC | 0 mM CTRL | 0 mM CTRL | B-DNA | G4 | 0.001 | 0.005 | 0.23 | 0.04 |
| cMYC | motif | 0 mM CTRL | 5 mM MnO<sub>4</sub> | B-DNA | B-DNA | 0.286 | 0.013 | 0.20 | 0.00 |
| cMYC | RC | 0 mM CTRL | 5 mM MnO<sub>4</sub> | B-DNA | B-DNA | 0.044 | 0.074 | 0.15 | 0.03 |
| cMYC | motif | 0 mM CTRL | 5 mM MnO<sub>4</sub> | G4 | G4 | 0.152 | 0.090 | 0.43 | 0.08 |
| cMYC | RC | 0 mM CTRL | 5 mM MnO<sub>4</sub> | G4 | G4 | 0.001 | 0.006 | 0.25 | 0.04 |
| VEGF-NMR | motif | 0 mM CTRL | 5 mM MnO<sub>4</sub> | B-DNA | B-DNA | 0.204 | 0.263 | 0.17 | 0.07 |
| VEGF-NMR | RC | 0 mM CTRL | 5 mM MnO<sub>4</sub> | B-DNA | B-DNA | 0.298 | 0.287 | 0.12 | 0.04 |
| VEGF-NMR | motif | 0 mM CTRL | 5 mM MnO<sub>4</sub> | G4 | G4 | 0.256 | 0.254 | 0.32 | 0.13 |
| VEGF-NMR | RC | 0 mM CTRL | 5 mM MnO<sub>4</sub> | G4 | G4 | 0.000 | 0.000 | 0.41 | 0.08 |
| VEGF-WT | motif | 0 mM CTRL | 5 mM MnO<sub>4</sub> | B-DNA | B-DNA | 0.157 | 0.197 | 0.19 | 0.08 |
| VEGF-WT | RC | 0 mM CTRL | 5 mM MnO<sub>4</sub> | B-DNA | B-DNA | 0.041 | 0.149 | 0.15 | 0.05 |
| VEGF-WT | motif | 0 mM CTRL | 5 mM MnO<sub>4</sub> | G4 | G4 | 0.257 | 0.218 | 0.26 | 0.07 |
| VEGF-WT | RC | 0 mM CTRL | 5 mM MnO<sub>4</sub> | G4 | G4 | 0.000 | 0.005 | 0.31 | 0.06 |

a: CTRL = control with no permanganate treatment. MnO<sub>4</sub> = KMnO<sub>4</sub> = potassium permanganate.
b: *p* = 0.000 indicates that an effect size greater than the experiment was not observed in 1000 permutations.
c: sd = Standard deviation that reflects test sensitivity to the read composition.
d: The test statistic is the sum of squared differences in mean DTW current error at each nt position.

We also investigated the composition of the ONT DNA 10.4.1 sequencing reaction and G4 structure stability in these reagents. The ONT sequencing reaction is composed of three proprietary components: elution buffer (EB), library beads (LIB) suspended in a buffer called library solution (LIS), and sequencing buffer (SB). These reagents are mixed at a ratio of 32:68:100 (Figure S4A). Elution buffer is typically a 10 mM Tris-HCl buffer (pH 8 to 9) containing a low concentration of detergent and has negligible UV absorbance. LIS also has negligible UV absorbance (Figure S4B) and supports a G4 CD signature at a 1x concentration (Figure S4C). In contrast, SB has extreme UV absorbance at 260 nm, making direct CD detection of G4 structure difficult (Figure S4B). Thus, the ONT sequencing reaction has at least one component that supports G4 structure.

### Machine learning identifies original structure state based on nanopore signal

The large difference between B-DNA and G4 DTW current error at the same motif indicates that DTW current error should be an effective classifier for the original structure state. We therefore combined our data from all three G4 motifs, aligned the datasets at the first nt of the G4 motif, and randomly split the data into a training set (146,366 reads) and a test set (36,592 reads). We then trained four model architectures—logistic regression, random forest, XGBoost, and a 1D Convolutional Neural Network (CNN)—to classify reads originating from B-DNA or G4 structure (Figure 2A). All four architectures had excellent performance on the test set, with the XGBoost algorithm having the highest area under the curve (AUC; Figure 2B; max AUC = 0.966). The classifier worked equally well on all motifs (Figure 2C), where the false positive rate (4%) was lower than the false negative rate (14%). Interestingly, the most important nt in the classifier occurred 12 nt upstream of the start of the G4 motif, approximately the distance at which the G4 structure would begin interacting with the pore as the DNA strand translocates through the nanopore membrane^19^.

**Figure 2.**
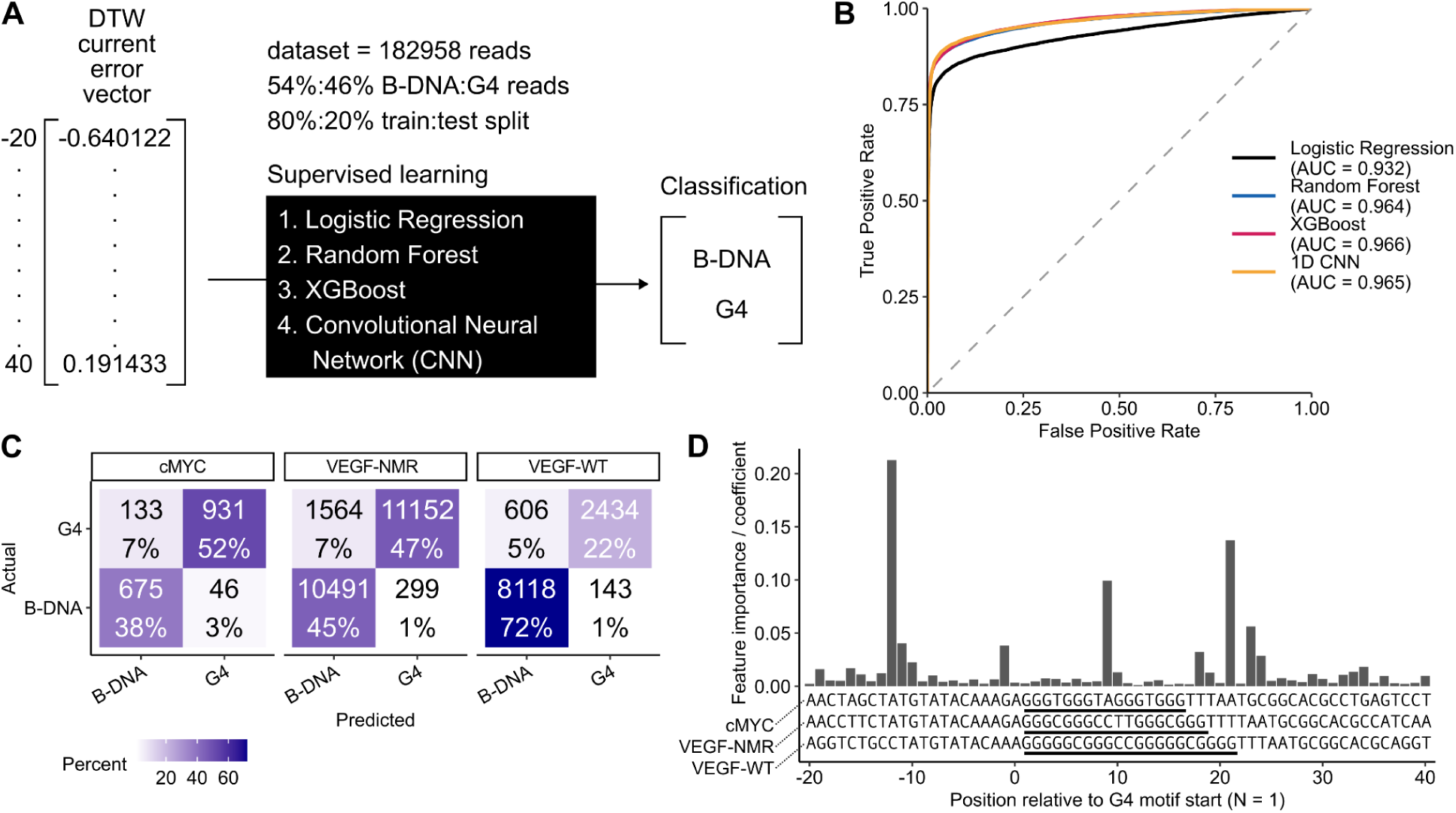
Machine learning accurately predicts structure state using the DTW current error (pA). **(A)** ML training and testing experiment design. **(B)** Receiver operating characteristic curves comparing model architecture performance. Area under the curve (AUC) values evaluate classification, where values of 1.0 and 0.5 indicate perfect and random classification, respectively. **(C)** Confusion matrices constructed on the test set to evaluate true positive, false positive, true negative, and false negative rates for the XGBoost model. The values represent the number of reads. The denominator in the percent calculation is the number of reads originating from each G4 motif. **(D)** Feature importance coefficients calculated using the average gain in the XGBoost loss function attributed to each nt.

### Chemical probing with permanganate followed by direct ONT sequencing identifies single-stranded DNA in vitro

While accurate on carefully prepared *in vitro* models, direct detection of G4 structure from whole-genome sequencing data is challenging (because a controlled experiment with G4 folding is difficult to design in the cell), and the structure may not be retained during DNA extraction and library preparation. Chemical probes that react rapidly with DNA in the cell, thereby locking secondary structure information into the chemical signature of the DNA fragment, have the potential to provide a link between the sequencing signal and the original secondary structure state (Figure 3A). We used potassium permanganate as a starting point because it specifically modifies single-stranded Ts and Cs, thus providing information on a wide range of potential alternative structures^20–23^. Also, potassium permanganate is relatively safe to handle compared with other nucleic acid-modifying reagents.

**Figure 3.**
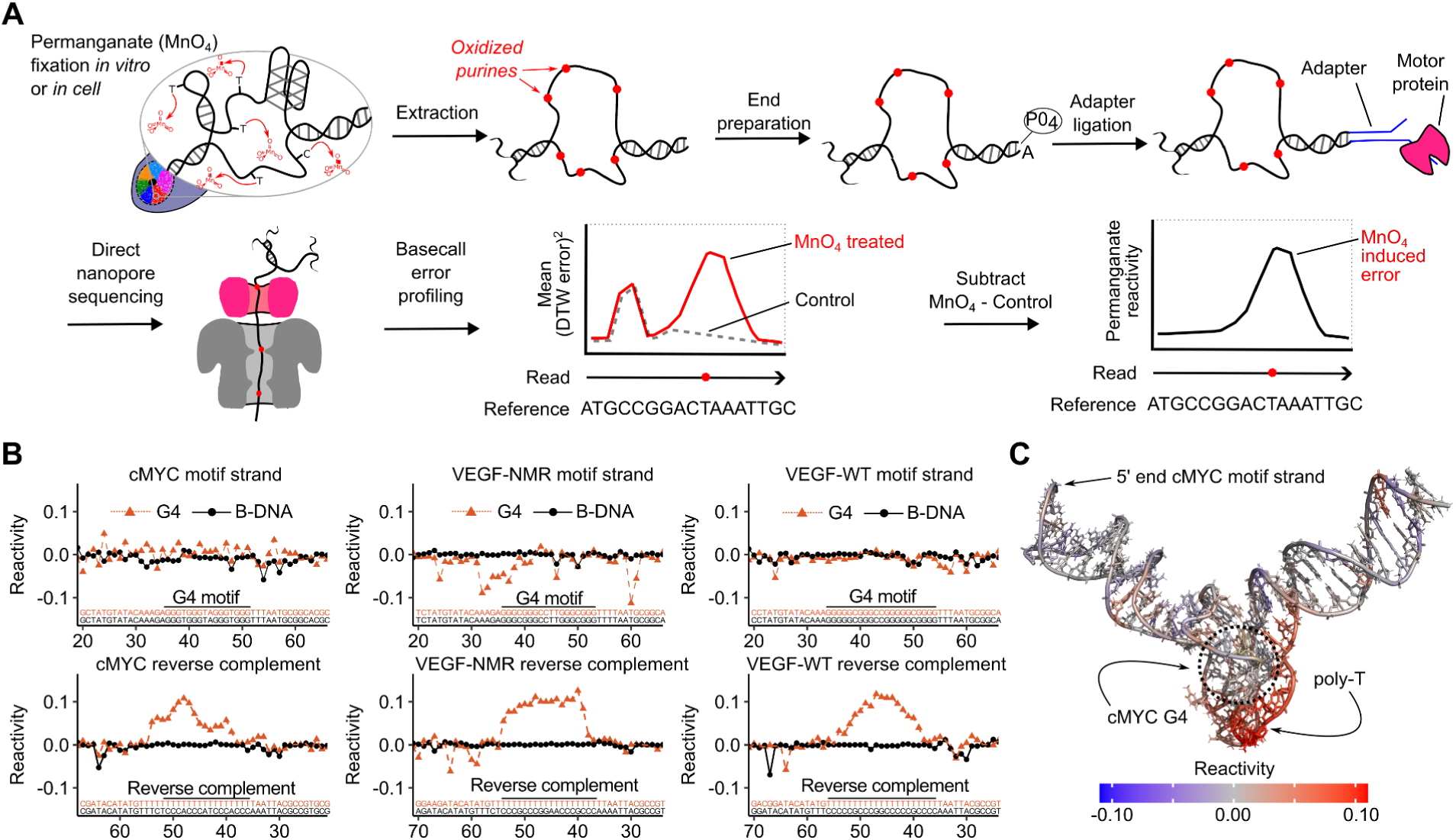
Permanganate footprinting with direct ONT sequencing identifies ssDNA locus *in vitro.* **(A)** Library preparation and data analysis for detecting permanganate reactive loci. **(B)** Permanganate reactivity at each nt for G4-duplex models. Reactivity was calculated as the difference between the mean squared DTW current error in the treatment and control. **(C)** Permanganate reactivity mapped onto the 3D structure of the *cMYC* G4-duplex model (Protein Databank ID 8DUT) using Pymol (Schrodinger).

As a proof of concept, we treated the G4-duplex models with potassium permanganate *in vitro* and constructed DTW current error profiles from the resulting ONT reads (Table S1, Figure S5-S7). Using the *cMYC* G4-duplex model as an example, permanganate treatment did not cause a significantly different DTW current error profile for the G4 motif strand in the G4 or B-DNA conformation (Table 2 Functional permutation test ^18^; *p* = 0.15±0.09 or *p* = 0.28±0.01, respectively permutation test with subsampling^18^ see Methods for details). In contrast, permanganate did cause a significantly different DTW current error for the RC strand in the G4 but not the B-DNA conformation (Table 2; *p* = 0.001±0.006 and *p* = 0.04±0.07, respectively), consistent with the former strand being in the single-stranded state.

We then calculated permanganate reactivity for each G4-duplex model, where the squared mean DTW current error of the control was subtracted from that for the treated sample at each nt (Figure 3A). Permanganate reactivity was near 0 for G4-duplex models in the B-DNA conformation (Figure 3B). However, permanganate reactivity exhibited a prominent increase on the single-stranded poly-T RC for every G4-duplex model in the G4 conformation. Likewise, the regions of high permanganate reactivity mapped to solvent-exposed Ts on the three-dimensional structure (Figure 3C).

Interestingly, permanganate reactivity was more stochastic on the motif strand in the G4 structure than the B-DNA structure (Figure 3B), often producing negative values. This signal is unlikely to be noise alone, given the massive sequencing depth of these experiments, and likely reflects uncharacterized interactions between the secondary structure, the permanganate modifications, and the nanopores.

### Chemical probing with permanganate followed by direct ONT sequencing identifies single-stranded DNA in cells

We performed permanganate footprinting in a human lymphoblastoid cell line—on two human, HG002-GM24385 cell-line culture replicates. Each culture replicate was split in half, and treated with either no (0 mM) or 20 mM permanganate. DNA was then extracted and sequenced to final approximate coverage of ∼30× and ∼15× for the 0 mM and 20 mM samples, respectively (Table S3). Reads from the 0 mM samples were slightly longer than reads from the 20 mM samples, 12-13 kbp and 10-11 kbp, respectively (Figure 4A). However, the final alignment accuracy was very high, with almost all the sequencing reads aligning with a near-perfect MAPQ of 60 (Figure 4B).

**Figure 4.**
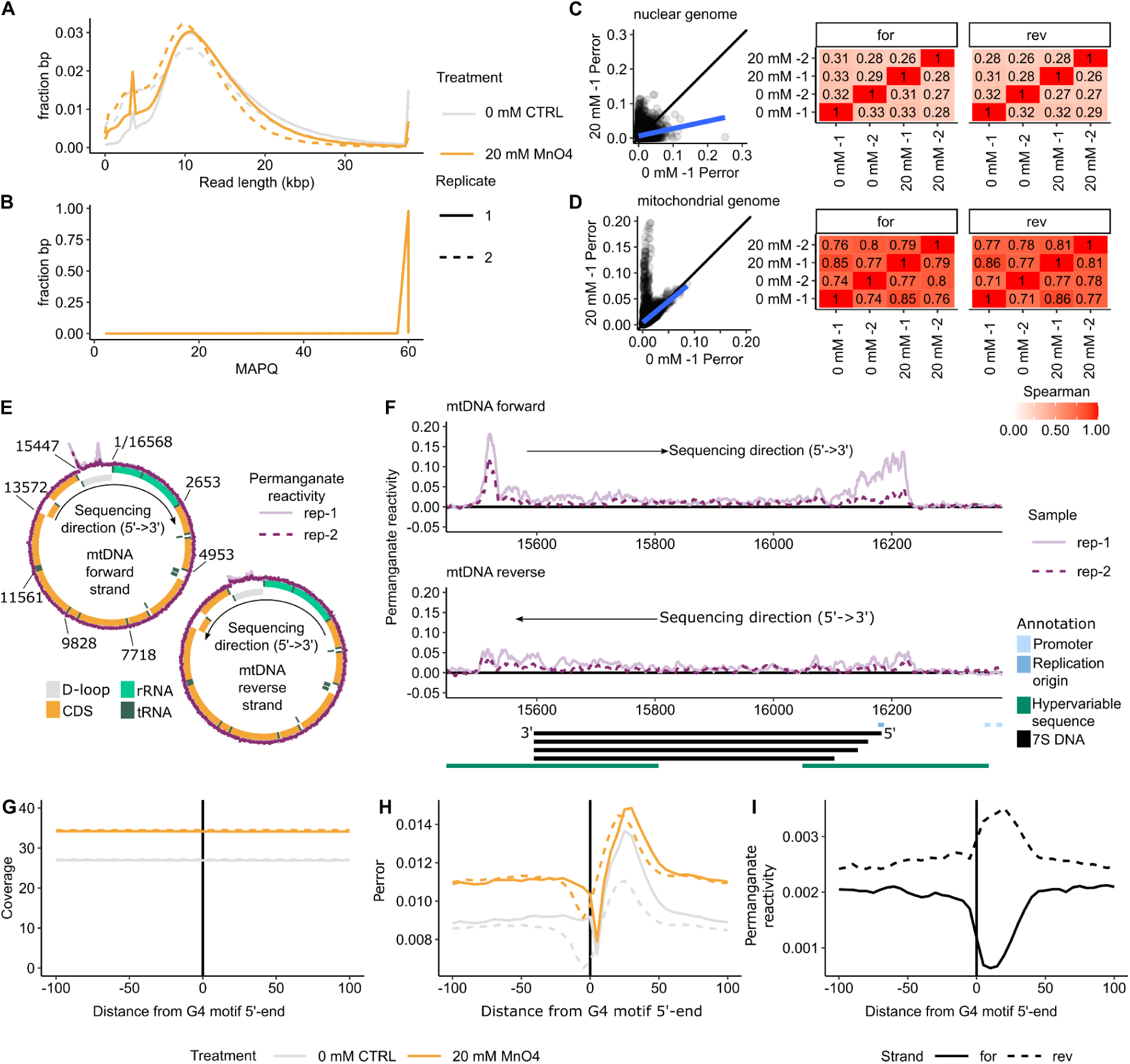
Permanganate footprinting with direct ONT sequencing identifies ssDNA *in cells.* Two experiments where a culture of the HG002-GM24385 cell line was split and either treated with water or 20 mM potassium permanganate are shown. **(A)** Read length histograms by sample. Y-axis are proportional to base-pairs sequenced instead of number of reads. The peak at 3.6 kb corresponds to an amplicon of the Lambda phage genome included as a spike in control. **(B)** Mapping quality (MAPQ) histograms. MAPQ is calculated with Minimap2 and corresponds to MAPQ = -10*log10(P_mapping error_) where P_mapping error_ is the probability that the alignment position is wrong. MAPQ = 60 is the highest value reported by Minimap2. Curves from different samples are superimposable at this scale. **(C)** Permanganate footprinting error correlation between samples across the nuclear genome. (Left): P_error_ in the 20 mM-1 sample versus 0 mM -1 sample from 10,000 randomly selected nt on the forward strand and reverse strand (20,000 nt total). The black line represents y = x and the blue line represents a best y = mx +b fit. (Right): Spearman correlation coefficients comparing all samples. **(D)** Same as panel C, but for the mitochondrial genome which was sequenced to a higher coverage. **(E)** Permanganate reactivity across the mitochondrial (mt) DNA genome. Two reactivity plots are shown, one for the forward (heavy) strand and one for the reverse (strand). The purple outer circle, middle circle, and inner circle correspond to the reactivity, forward annotations, and reverse annotations, respectively. **(F)** Zoomed in reactivity across the non-coding D-loop region of the mt genome. Plots share an x-axis with annotations, including binding sites for 4 known 7S DNA products. **(G)** Average coverage around G4 motifs on chromosome 1 aligned to the first nt of G4 motifs. Datasets were combined, coverage was smoothed in 5 nt windows, and reported as the mean across all G4s. **(H)** and **(I)** same plots as panel G for P_error_ and permanganate reactivity, respectively.

P_error_ rates were calculated across the human nuclear and mitochondrial (mt) genomes. DTW current errors were not calculated because of the high computational cost associated with raw nanopore signal alignment to the 3.1 Gbp human genome at 15×-30× coverage. Moreover, permanganate reactivity calculated with P_error_ was similar to permanganate reactivity calculated with DTW current errors (Figure S8). For the nuclear genome, P_error_ was poorly correlated between samples at the single-nt level (Figure 4C, Spearman’s Rho = 0.26-0.33). This poor correlation was expected because the single-nt coverage was reduced to 7.5× to 15× (15× to 30×/2 strands), likely too low to adequately capture the actual variance in P_error_. In contrast, the correlation was much higher for the mt genome (Figure 4D, Spearman’s Rho = 0.71-0.86), which is present at two orders of magnitude higher copy number in most somatic cells compared with that for the nuclear genome ^24^ and had a mean coverage of 610× (standard deviation = 206×) per strand in our samples.

Permanganate reactivity calculated using P_error_ was high at the non-coding, D-loop region of the mt genome in both replicates (Figure 4E). Permanganate reactivity was highest at both ends of the 7S DNA binding site on the forward (heavy) strand (Figure 4F). Permanganate reactivity was also elevated on the reverse (light) strand with no large peaks. Thus, permanganate reactive mtDNA regions in the cell correspond to loci where elevated rates of ssDNA are expected. Indeed, the region on mtDNA corresponding to the 7S DNA binding site forms a unique three-stranded region, including ssDNA, also called the displacement loop (D-loop) ^25^.

Last, we searched for the permanganate reactivity patterns we observed for our G4-duplex models *in vitro* at G4 motifs in the cell. To overcome noise from low coverage of the nuclear genome, we combined sequencing replicates, recalculated coverage, P_error,_ and permanganate reactivity, performed a 5-nt smooth, and averaged across all G4 motifs on chromosome 1 (Figure 4G-I). While this coverage did not allow one to identify structures at a single locus, we expected it to confirm the pattern we observed *in vitro* (Figure 3B). G4 motifs had no effect on coverage for both strands of the 0 mM and 20 mM samples (Figure 4G). P_error_ was elevated in 20 mM samples compared with 0 mM samples and peaked at ∼25 N from the first nt of G4 motifs, consistent with permanganate modification and known genome-wide sequencing errors, respectively (Figure 4H). Last, as expected, permanganate reactivity was higher at G4 motifs on the reverse strand (Figure 4I), consistent with *in vitro* models (Figure 3B). This pattern was confirmed on chromosomes 9 and 19 (Figure S9).

## Discussion

We found that DNA secondary structure and modifications introduced by a chemical probe can be detected in direct ONT sequencing data. While both signals represent promising approaches to studying alternative, non-B DNA secondary structures, our experiments identify methodological constraints that must be overcome to achieve high resolution and specificity for this purpose in large genomes.

Sequencing biases at G4 motifs are well established^4^. Indeed, chemically induced G4 structures in whole-genome Illumina sequencing runs were previously used to identify sequence motifs capable of forming such structures^26^. Using G4-duplex models, we observed that raw nanopore current signal at G4 motifs is dependent on DNA conformation (Figure 1). Furthermore, machine learning can identify the original conformation at the single-molecule resolution using the same current signal (Figure 2). This observation implies that one could also screen whole-genome ONT sequencing reads for loci in the G4 conformations. However, our present *in vitro* dataset is limited to three closely related G4 motifs, and it is unknown how our observations will generalize to a more diverse array of motifs. More work is also needed to establish the persistence of alternative secondary structures in ONT library preparations. Also, researchers should consider that the data this strategy would be based on, raw ONT current signal, is not available in the National Center for Biotechnology Information Sequence Read Archive or saved by default in current versions of ONT data collection software—something that we suggest should be recorded on par with interpulse duration (IPD) data for Pacific Biosciences data.

Chemical reactivity is an established readout of *in cell* RNA secondary structures^27,28^ and has been adapted to long-read sequencing^29–32^, although these strategies usually rely on reverse-transcriptase errors to convert chemical structure to a sequence-readable signal. SsDNA associated with R-loops has also been identified at single-molecule resolution using chemical probing and long-read-sequencing^33^; this approach relies on cytosine to uracil conversion with sodium bisulfite after DNA extraction. Extensive work has established chemical probing of DNA structures using short-read protocols^21,22,34,35^. Adapting antibody-based non-B DNA detection techniques to long reads is also a promising alternative strategy ^36^. Herein, we demonstrated that permanganate footprinting with direct nanopore sequencing can identify ssDNA associated with G4 folding *in vitro* and *in cell* (Figure 3-4). The primary experimental consideration is the size and copy number of the region of interest, where high coverage, resolution, and specificity were achieved for short G4-duplex models *in vitro* and for the mt genome *in cell*. However, useful coverage, resolution, and specificity were not achieved for the entire human genome. Higher coverage, improved chemical probing conditions, and sensitive footprint detection strategies are expected to alleviate this limitation in the future.

## Methods

### G4-duplex model DNA preparation

Model DNA oligos (Table 1) were ordered from Integrated DNA Technologies (IDT), resuspended in nuclease-free water to a target final 100 µM concentration, and purified with denaturing polyacrylamide gel (PAGE) electrophoresis. Prior to loading, samples were heated to 95°C for 1.5 min at a final concentration of 50 µM DNA, 0.05×Tris borate ethylenediaminetetraacetic acid (EDTA) buffer (TBE), 0.0125% bromophenol blue, 16.7 mM EDTA pH 8.0, and 45.5% (v/v) formamide. Hot samples were immediately fractionated on a preheated 10% PAGE gel containing 1xTBE and 8.3 M urea. Full length DNA bands were identified by a near-UV shadow on a fluorescent TLC plate, excised, pulverized, and soaked overnight (rocking at room temperature) in 6 mL of 0.3 M sodium acetate 1 mM EDTA pH 8.0 buffer per gram gel fragment. The buffer now containing DNA fragments was passed through a 0.2 µm filter to remove gel fragments, mixed with a 3× volume of ice cold absolute ethanol, and incubated at -20°C for at least 2 hours. DNA was precipitated at 2,000 ×g for 30 min and dried at room temperature for 1 hour. Samples were resuspended, then buffer exchanged into 140 mM LiCl, 20 mM 3-(Morpholin-4-yl)propane-1-sulfonic acid (MOPS)-LiOH pH 7.0 buffer (total exchange volume > 1000:1) using 0.5 mL Amicon Ultra Centrifugal filters (Millipore) of the appropriate molecular weight cutoff (3 or 10 kDa). DNA duplexes were prepared by mixing equimolar concentrations of motif-containing strands with RC strands in a background of 140 mM LiCl, 20 mM MOPS-LiOH pH 7.0 buffer. Samples were heated to 95°C, slowly cooled to room temperature over the course of an hour, and stored at 4°C. Duplex DNA was incubated in 100 mM KCl, 140 mM LiCl, 20 mM MOPS-LiOH pH 7.0 buffer at 37°C for 1 hour to stabilize G4 structure immediately prior to CD, native gel electrophoresis, permanganate treatment, or library preparation.

### Circular dichroism (CD)

CD spectra were collected in quartz cuvettes at 25°C on a Jasco J-1500 spectrometer with the following settings: photometric mode-CD; HT. Abs, CD scale: 200 mdeg/1.0 d0D; FL scale: 200 mdeg/1.0 d0D; DIT: 4 sec; bandwidth: 1 nm; scanning speed: 50 nm/min; baseline correction: No Correction; accumulation: 3 times. The sample chamber and lamp were continuously purged with dry nitrogen to prevent condensation of ambient water. Buffer CD was subtracted at each wavelength. Molar ellipticity (*e*) was calculated using the following equation:

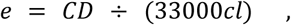

where *CD* is the circular dichroism in mDeg, *c* is the precise molar concentration, and *l* is the path length in cm. Concentrations are specified in Figure legends.

### Native gel electrophoresis

Five pmol of DNA were diluted to a final volume of 10 µL in the buffer it was folded in. Samples were weighted by mixing in 2 µL of 60% (w/v) glycerol 1×tris acetate EDTA (TAE) buffer, 0.025% (w/v) bromophenol blue loading solution. Samples were fractionated at 4°C on a 10% PAGE gel containing 1×TBE supplemented with 10 mM KCl. The gel was stained with 1×SYBR gold working reagent and visualized on a BioRad Gel Doc Go imager.

### Permanganate footprinting in vitro

G4-duplex models were multiplexed to a final concentration of 1 µM each model (3 µM total) in a final volume of 20 µL. Samples were quickly mixed with 6.6 µL of aqueous 5 mM potassium permanganate, incubated for exactly 80 seconds, and quenched with 26.6 µL of 100 mM β-mercaptoethanol 50 mM EDTA buffer.

### Sequencing of model G4-motifs

G4-duplex models were sequenced using an ONT Ligation Sequencing Kit V14 (Cat# SQK-LSK114) and companion reagents from New England Biolabs (NEB) with key adaptations for the unusually short sequence length. Samples were cleaned with a Zymo DNA Clean & Concentrator kit according to the manufacturer’s instructions using 5 volumes of DNA Binding Buffer and eluting with 50 μL nuclease-free water for 5 minutes. End-preparation reactions were prepared by mixing 48 µL DNA sample, 7 µL FFPE DNA Repair Buffer v2, 2 µL FFPE DNA Repair Mix, and 3 µL of Ultra II end-prep enzyme mix from a NEBNext Companion Module v2 for ONT (NEB Cat# E7672). End-preparation reactions were incubated at 20°C for 5 min and heat-inactivated at 65°C for 5 min. End-preparation reactions were cleaned with a Zymo DNA Clean & Concentrator kit according to the manufacturer’s instructions using 5 volumes of DNA Binding Buffer and eluting with 32 μL. Then, ligation reactions were prepared by mixing 30 µL of end-prepared DNA, 2.5 µL of ONT ligation adapter, 12.5 µL of ONT ligation buffer, and 5 µL of Salt-T4 DNA ligase from a NEBNext Companion Module v2 for ONT (NEB Cat# E7672). Ligation reactions were incubated for 10 min at room temperature, and cleaned with 40 µL AMPure XP beads provided by ONT and 250 µL ONT short fragment buffer. Library products were eluted in 35 µL ONT elution buffer and prepared for sequencing according to ligation sequencing kit instructions. Samples were sequenced using an ONT PromethION2 integrated (P2i) instrument and PromethION flow cells (Cat# FLO-PRO114M) until approximately 200,000 reads accumulated. Flow cells were washed with ONT flow cell wash kits (Cat# EXP-WSH004) between runs.

### Cell culture

HG002-GM24385 cells (Coriell Institute for Medical Research) were cultured in Roswell Park Memorial Institute (RPMI) medium 1640 (1×) with L-glutamine (Gibco) supplemented with 15% fetal bovine serum (FBS; Gemini Bio), 1×Penicillin Streptomycin Solution (Pen/Strep; Corning), and freshly added 1× GlutaMAX (Gibco) in a 5% CO_2_ incubator at 37°C. The protocols for handling, storage, subculturing, and cryopreservation are available on the manufacturer’s website. Cells were grown to ∼1 million cells/mL prior to harvesting and treatment.

### Permanganate footprinting in cells

Five million cells were washed twice with ∼10 mL of pre-warmed 37°C Dulbecco’s phosphate buffered saline (DPBS) and resuspended in 2 mL of pre-warmed 15 mM Tris-HCl pH 7.5, 15 mM NaCl, 60 mM KCl, 300 mM Sucrose, 0.5 mM EGTA buffer for 60 seconds at room temperature. Cells were treated by adding 667 µL of either prewarmed water (Control) or potassium permanganate (20 mM) working solutions. A timer was started on the addition of the treatment solution, while samples were covered, gently mixed, and maintained at 37°C. The treatment solution was quenched and cells were lysed at exactly 80 seconds by the addition of 2.67 mL of 50 mM EDTA pH 8.0, 700 mM beta-Mercaptoethanol, 1% SDS. The lysate was then digested with 300 µg/mL proteinase K at 37°C for ∼20 hours (overnight).

DNA was extracted by adding 6 mL of phenol:chloroform:isoamyl alcohol (25:24:1) to the lysate. Samples were centrifuged at 2,000 ×g for 5 min to separate phases. The supernatant was transferred to a new 50 mL tube, then mixed with 4 mL of 5M ammonium acetate (final concentration of 2M) and 20 mL of ice-cold absolute ethanol. DNA was further incubated at -20°C for >30 min and precipitated for 60 minutes at 4°C, 2,000 ×g. The pellet was washed with 2 mL ice-cold 80% ethanol, then reprecipitated for 5 minutes at 4°C, 2,000 ×g. The pellet was dried for 60 minutes and resuspended in 1 mL of 10 mM Tris-HCl pH 8.0 at room temperature overnight.

RNA was removed by adding 2.5 μL of 20 mg/mL RNase A and incubating at 37°C for 1 hour. DNA was extracted by adding 1 mL of phenol:chloroform:isoamyl alcohol (25:24:1). Samples were spun at 2,000 ×g for 5 min to separate phases. The supernatant was transferred to a new 15 mL tube, then mixed with 0.667 mL of 5M ammonium acetate (final concentration of 2M) and 3.34 mL of ice-cold absolute ethanol. DNA was further incubated at -20°C for >30 min and then precipitated by centrifugation for 60 minutes at 4°C, 2,000 ×g. The pellet was dislodged in 1 mL ice-cold 80% ethanol and then reprecipitated for 5 minutes at 4°C, 2,000 ×g. The pellet was dried for 60 min and resuspended in 1.0 mL of 10 mM Tris-HCl pH 8.0. Samples were incubated at room temperature for ∼20 hours (overnight) and then stored at 4°C (for ∼3 days) until the DNA solution was homogenous. Homogeneity was assessed by measuring concentrations of the top, middle, and bottom of the solution using a Nanodrop microvolume spectrometer (Thermo Fisher Scientific) and Qubit HS dsDNA assay (Invitrogen/Thermo Fisher).

### Sequencing of in-cell permanganate-treated DNA

Sequencing libraries were prepared using 2 µg of genomic DNA. First, the DNA was sheared to ∼10-15 kbp using a Covaris g-TUBE (Cat# 520104). Libraries were then prepared using an ONT Ligation Sequencing Kit V14 (Cat# SQK-LSK114) according to the manufacturer’s instructions. Final library concentrations were determined with a Qubit HS dsDNA assay, and 500 ng of library was sequenced using an ONT P2i instrument on new PromethION flow cells (Cat# FLO-PRO114M) for ∼48 hours. Flow cells were washed with ONT flow cell wash kits (Cat# EXP-WSH004) before loading and sequencing another 500 ng of library until negligible pore activity was collected.

### Nanopore secondary structure and chemical footprinting data analysis

Raw nanopore sequencing data were processed using the NanoPrint toolkit https://github.com/makovalab-psu/NanoPrint_toolkit, a Snakemake pipeline for strand-resolved, per-base secondary structure and chemical footprinting analysis of ONT long-read data. Raw electrical signal data (pod5 format) were basecalled using Dorado (ONT; Dorado version 1.1.1+3c7eef9+cu12080, libtorch version 2.6.0) with the super-accuracy model (DNA 10.4.1 e8.2 400bps sup v5.2.0). Dorado was used with the “--emit-moves” flag to retain move tables for downstream signal alignment. Basecalled reads were aligned to reference sequences using minimap2 (version 2.28-r1209)^37^ with the long-read high-quality preset (lr:hq), preserving all move table tags in the output BAM. Alignments were filtered to retain only primary alignments with mapping quality ≥ 20. Per-base error (P_error_) was calculated for each strand (forward/reverse) by taking the mean base call error probability after transforming Phred scores to a probability (P_error_) using the following equation:

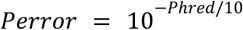

Filtered alignments were re-processed with Uncalled4 (version 4.1.0)^17^ to perform dynamic time warping (DTW) alignment of the raw ionic current signal against the expected pore model signal. Pore model alignment was performed with “uncalled4 align” with the “--min-aln-length 50” flag to accommodate short G4-duplex models. The dtw.current (pA), dtw.current_sd (pA), dtw.start (events), dtw.length (events), and dtw.model_diff (pA; called DTW current error elsewhere) were extracted to a tab-delimited text format for each aligned position using “uncalled4 convert”. Chemical reactivity (pA^2^) was computed as the difference between the treatment and control mean squared DTW current error (Σdtw.model_diff²/N). Dependencies are specified on GitHub and can be installed using Conda or Mamba. Alternatively, reactivity was computed as the difference between the treatment and control P_error_.

G4-duplex model data were mapped to the sequences in Table 1, retaining only forward strand alignments. *In cell* HG002-GM24385 permanganate footprinting data were mapped to the paternal autosome haplotypes, the sex chromosomes, and the mitochondrial genome from the HG002 Q100 project ^38^. Mitochondrial annotations from NCBI Reference Sequence NC_012920.1 ^39^ were converted to HG002 coordinates using Liftoff ^40^. Noncoding, D-loop region annotations were compiled from ^25^. G4 annotations were provided by the G4Discovery pipeline ^41^.

### Permutation test for comparing DTW current error profiles between conditions

We used a repeated subsampling permutation test to compare DTW signal error profiles (Table S1) ^18^. This test calculates the probability that two conditions have the same average DTW current error profile while accounting for effects from the extremely large sample size. Intuitively, this tests whether a reader could easily distinguish two 100-read DTW current error (pA) matrices shown in Figure 1B-E. The procedure was as follows. First, 100 reads were drawn randomly from each condition, and their per-position DTW current error values were arranged into a DTW current error-by-position matrix.

Subsampling was used instead of testing all reads simultaneously because the large number of sequenced molecules renders the test sensitive to trivial deviations. Second, the sum of squared differences in mean DTW current error at each nt position was calculated to generate a test statistic, and corresponds to the effect size in pA^2^. Third, a null distribution for this statistic was constructed by randomly permuting read assignments between the two groups 1,000 times and recomputing the statistic. Fourth, the p-value was calculated as the proportion of permuted statistics greater than or equal to the observed value. This process was repeated 100 times. The median and standard deviation of the P-value distribution are reported in Table 2 and Table S2.

This procedure was modified for the VEGF models when DTW current error profiles were compared between the perfect and poly-T RC strands. The VEGF perfect RC is four nt longer than the poly-T RC, creating a misalignment of flanking sequences that inflates variation between samples. Thus, during construction of the reads-by-position matrix, both flanking sequences were aligned, and 3 random nt were dropped from the RC of the G4 motif at every subsample.

### Secondary structure machine learning training and testing

Machine learning was performed using Python 3.11.15, and random seeds were fixed at 42. Per-read feature vectors composed of DTW current error were extracted with “uncalled4 convert” then converted to a G4-relative coordinate system where N = 1 corresponds to the first nt of the G4 motif. Positions less than -20 and greater than +40 were removed to avoid contamination with spurious helix-end effects, resulting in a total of 61 nt. Reads with more than 30 missing nt and more than 5 consecutive missing nt were discarded. Missing nt gaps of less than 5 nt were filled with the average of two adjacent non-missing nts, and leading/trailing missing nts were filled by extension of the nearest observed nt. Reads from all three G4 motifs were combined, retaining motif identity and original structure labels. An 80:20 stratified train/test split (meaning that the 80/20 ratio was enforced within subgroups = 3 motifs × 2 structure classes) was performed using the scikit-learn train_test_split function (version 1.9.0).

Logistic Regression was trained with scikit-learn using an L2 regularization penalty (C = 1.0) and a maximum of 1,000 solver iterations. A constant-zero imputer was included to prevent errors from residual missing values. Random Forest was trained with scikit-learn using 100 decision trees, allowing full tree growth. eXtreme Gradient Boosting (XGBoost; version 3.2.0) was trained with 100 boosting rounds, a maximum tree depth of 6, and a learning rate of 0.1, using log-loss as the evaluation metric. Feature importances correspond to gain-based scores, which represent the average gain in the loss function attributed to each nt. The 1D Convolutional Neural Network (CNN) was implemented in PyTorch (version 2.12.1) and was composed of two convolutional blocks. The first block had 32 filters of kernel size 5 (padding 2) followed by max-pooling. The second block had 64 filters of kernel size 3 (padding 1) followed by max-pooling. Thus, the 61-position input was reduced to 15 positions before flattening. The flattened representation was passed through a fully connected layer of 128 units with ReLU activation and 50% dropout, followed by a single output unit. The model was trained with a batch size of 64 for 50 epochs using the Adam optimizer (learning rate 0.001) and binary cross-entropy with logit loss. Models were evaluated on the test set and evaluations included the receiver operating characteristic curve (AUROC), confusion matrices, and feature importance profile.

## Generative artificial intelligence (AI) contribution

Claude Code (Anthropic; Sonnet and Opus models) was used for data analysis programming focusing on pipeline construction, efficient resource utilization, formatting, documentation, and debugging. Statistical analysis and Figures were designed and performed by JPS with trivial AI contribution. The manuscript was drafted by JPS using Claude Code only to initially compile and cross validate descriptions of computational methods. Claude was also used to proofread the manuscript for typographical mistakes.

## Author contributions

This study was designed by JPS and KDM. Wet lab experiments were performed by JPS, HZ, and LH. Data analysis was performed by JPS and LH. The first draft of the manuscript was compiled by JPS and edited by KDM. Funding and resources were provided by KDM.

## Acknowledgments and Funding

We are grateful to Francesca Chiramonte for advising on statistical analysis and figures. We thank Byung June Ko for help with early versions of data analysis code to calculate P_error_ at scale. This work was supported by the NIH grant R35 GM151945 and by the Willaman Endowment Fund to KDM.

## Notes

### Competing Interest Statement

The authors have declared no competing interest.

## References

1. Makova, K. D. & Weissensteiner, M. H. Noncanonical DNA structures are drivers of genome evolution. Trends Genet. 39, 109–124 (2023).

2. Makova, K. D., Mohanty, S., Smeds, L., Sieg, J. & Lahnsteiner, A. Unraveling non-B DNA structures in the era of telomere-to-telomere genomes. Annu. Rev. Genomics Hum. Genet. (2026) doi:10.1146/annurev-genom-120324-010100.

3. Morrissey, A., Shi, J., James, D. Q. & Mahony, S. Accurate allocation of multimapped reads enables regulatory element analysis at repeats. Genome Res. 34, 937–951 (2024).

4. Weissensteiner, M. H. et al. Accurate sequencing of DNA motifs able to form alternative (non-B) structures. Genome Res. 33, 907–922 (2023).

5. Li, H. & Durbin, R. Genome assembly in the telomere-to-telomere era. Nat. Rev. Genet. 25, 658–670 (2024).

6. Formenti, G., et al. The complete genome of a songbird. bioRxivorg (2025) doi:10.1101/2025.10.14.682431.

7. Smeds, L., et al. Non-canonical DNA in bird telomere-to-telomere genomes. bioRxivorg (2025) doi:10.1101/2025.10.17.683159.

8. Stergachis, A. B., Debo, B. M., Haugen, E., Churchman, L. S. & Stamatoyannopoulos, J. A. Single-molecule regulatory architectures captured by chromatin fiber sequencing. Science 368, 1449–1454 (2020).

9. Altemose, N. et al. DiMeLo-seq: a long-read, single-molecule method for mapping protein–DNA interactions genome wide. Nat. Methods 19, 711–723 (2022).

10. Fu, Y., Timp, W. & Sedlazeck, F. J. Computational analysis of DNA methylation from long-read sequencing. Nat. Rev. Genet. 26, 620–634 (2025).

11. Guiblet, W. M. et al. Long-read sequencing technology indicates genome-wide effects of non-B DNA on polymerization speed and error rate. Genome Res. 28, 1767–1778 (2018).

12. Hosseini, M. et al. Deep statistical modelling of nanopore sequencing translocation times reveals latent non-B DNA structures. Bioinformatics 39, i242–i251 (2023).

13. Monsen, R. C., Chua, E. Y. D., Hopkins, J. B., Chaires, J. B. & Trent, J. O. Structure of a 28.5 kDa duplex-embedded G-quadruplex system resolved to 7.4 Å resolution with cryo-EM. Nucleic Acids Res. 51, 1943–1959 (2023).

14. Fleming, A. M., Guerra Castañaza Jenkins, B. L., Buck, B. A. & Burrows, C. J. DNA Damage Accelerates G-Quadruplex Folding in a Duplex-G-Quadruplex-Duplex Context. J. Am. Chem. Soc. 146, 11364–11370 (2024).

15. Ambrus, A., Chen, D., Dai, J., Jones, R. A. & Yang, D. Solution structure of the biologically relevant G-quadruplex element in the human c-MYC promoter. Implications for G-quadruplex stabilization. Biochemistry 44, 2048–2058 (2005).

16. Agrawal, P., Hatzakis, E., Guo, K., Carver, M. & Yang, D. Solution structure of the major G-quadruplex formed in the human VEGF promoter in K+: insights into loop interactions of the parallel G-quadruplexes. Nucleic Acids Res. 41, 10584–10592 (2013).

17. Kovaka, S. et al. Uncalled4 improves nanopore DNA and RNA modification detection via fast and accurate signal alignment. Nat. Methods 22, 681–691 (2025).

18. Cremona, M. A. et al. IWTomics: testing high-resolution sequence-based ‘Omics’ data at multiple locations and scales. Bioinformatics 34, 2289–2291 (2018).

19. Van der Verren, S. E. et al. A dual-constriction biological nanopore resolves homonucleotide sequences with high fidelity. Nat. Biotechnol. 38, 1415–1420 (2020).

20. Kouzine, F. et al. Global Regulation of Promoter Melting in Naive Lymphocytes. Cell 153, 988–999 (2013).

21. Kouzine, F. et al. Permanganate/S1 Nuclease Footprinting Reveals Non-B DNA Structures with Regulatory Potential across a Mammalian Genome. Cell Syst. 4, 344–356.e7 (2017).

22. Lahnsteiner, A. et al. In vivo detection of DNA secondary structures using permanganate/S1 footprinting with direct adapter ligation and sequencing (PDAL-Seq). Methods Enzymol. 695, 159–191 (2024).

23. Sieg, J. et al. Comparative analysis of single-stranded and non-canonical DNA formation in human and other ape cells with telomere-to-telomere genomes. bioRxivorg (2025) doi:10.1101/2025.11.03.686349.

24. Ding, J. et al. Assessing mitochondrial DNA variation and copy number in lymphocytes of ∼2,000 Sardinians using tailored sequencing analysis tools. PLoS Genet. 11, e1005306 (2015).

25. Nicholls, T. J. & Minczuk, M. In D-loop: 40 years of mitochondrial 7S DNA. Exp. Gerontol. 56, 175–181 (2014).

26. Sahakyan, A. B. et al. Machine learning model for sequence-driven DNA G-quadruplex formation. Sci. Rep. 7, 14535 (2017).

27. Ding, Y. et al. In vivo genome-wide profiling of RNA secondary structure reveals novel regulatory features. Nature 505, 696–700 (2014).

28. Rouskin, S., Zubradt, M., Washietl, S., Kellis, M. & Weissman, J. S. Genome-wide probing of RNA structure reveals active unfolding of mRNA structures in vivo. Nature 505, 701–705 (2014).

29. Yang, M. et al. In vivo single-molecule analysis reveals COOLAIR RNA structural diversity. Nature 609, 394–399 (2022).

30. Bohn, P., Gribling-Burrer, A.-S., Ambi, U. B. & Smyth, R. P. Nano-DMS-MaP allows isoform-specific RNA structure determination. Nat. Methods 20, 849–859 (2023).

31. Bizuayehu, T. T. et al. Long-read single-molecule RNA structure sequencing using nanopore. Nucleic Acids Res. 50, e120–e120 (2022).

32. Gribling-Burrer, A.-S., Bohn, P. & Smyth, R. P. Isoform-specific RNA structure determination using Nano-DMS-MaP. Nat. Protoc. 19, 1835–1865 (2024).

33. Malig, M., Hartono, S. R., Giafaglione, J. M., Sanz, L. A. & Chedin, F. Ultra-deep coverage single-molecule R-loop footprinting reveals principles of R-loop formation. J. Mol. Biol. 432, 2271–2288 (2020).

34. Wu, T., Lyu, R., You, Q. & He, C. Kethoxal-assisted single-stranded DNA sequencing captures global transcription dynamics and enhancer activity in situ. Nat. Methods 17, 515–523 (2020).

35. Lyu, R. et al. KAS-seq: genome-wide sequencing of single-stranded DNA by N3-kethoxal–assisted labeling. Nat. Protoc. 17, 402–420 (2022).

36. Lyu, J., Shao, R., Kwong Yung, P. Y. & Elsässer, S. J. Genome-wide mapping of G-quadruplex structures with CUT&Tag. Nucleic Acids Res. 50, e13–e13 (2022).

37. Li, H. New strategies to improve minimap2 alignment accuracy. Bioinformatics 37, 4572–4574 (2021).

38. Hansen, N. F., et al. A complete diploid human genome benchmark for personalized genomics. bioRxivorg (2025) doi:10.1101/2025.09.21.677443.

39. Anderson, S. et al. Sequence and organization of the human mitochondrial genome. Nature 290, 457–465 (1981).

40. Shumate, A. & Salzberg, S. L. Liftoff: accurate mapping of gene annotations. Bioinformatics 37, 1639–1643 (2021).

41. Mohanty, S. K., et al. Variation and selection at predicted G-quadruplexes across the human pangenome. bioRxivorg (2026) doi:10.64898/2026.06.18.733261.

